# Contributions of Gonadal Hormones in the Sex-Specific Organization of Context Fear Learning

**DOI:** 10.1101/2022.08.01.501766

**Authors:** Lorianna Colón, Eduardo Peru, Damian G. Zuloaga, Andrew M. Poulos

## Abstract

It is widely established that gonadal hormones are fundamental to modulating and organizing the sex-specific nature of reproductive-related behaviors. Recently we proposed that context fear conditioning may emerge in a sex-specific manner organized prior to the pubertal surge of gonadal hormones. Here we sought to determine the necessity of male and female gonadal hormones secreted at critical periods of development upon context fear learning. We tested the organizational hypothesis that perinatal and pubertal gonadal hormones play a permanent role in organizing contextual fear learning. We demonstrate that the life-long absence of gonadal hormones by perinatal orchiectomy (oRX) in males and ovariectomy (oVX) in females resulted in a reduction of CFC in adult males and an enhancement of CFC in adult females. In females, the gradual introduction of estrogen before conditioning partially rescued this effect. However, the decrease of CFC in adult males was not rescued by introducing testosterone before conditioning. Next, at a further point in development, preventing the pubertal surge of gonadal hormones by prepubertal oRX in males resulted in a reduction in adult CFC. In contrast, in females, prepubertal oVX did not alter adult CFC. However, the adult introduction of estrogen in prepubertal oVX rats reduced adult CFC. Lastly, the adult-specific deletion of gonadal hormones by adult oRX or oVX alone or replacement of testosterone or estrogen did not alter CFC. Consistent with our hypothesis, we provide initial evidence that gonadal hormones at early periods of development exert a vital role in the organization and development of CFC in male and female rats.

## Introduction

Sex differences in learning and memory have been primarily investigated and understood to result from the activational or modulatory actions of gonadal hormones (see Daniel & Dohanich, 2001; Korol, 2004; Luine, 2008; Taxier et al., 2020 for reviews). However, both estrogens and androgens, produced and secreted by the ovaries and testes, can organize the development of sexual dimorphisms in motivated behavior and their underlying brain circuits (Lenz & McCarthy, 2010; Schulz & Sisk, 2016; Oyola & Handa, 2017). Evidence for the organizational role of these hormones in learning has been sparse (Williams et al., 1990). Recently we found evidence that contextual fear learning develops in a sex-specific manner initiated before puberty, suggesting that gonadal hormones during the perinatal period may contribute to the organization and development of contextual fear learning (Colón et al., 2018). To date, only a handful of studies have examined the necessity of these hormones in fear-motivated learning (Anagnostaras et al., 1998; Morgan & Pfaff, 2001; Gupta et al., 2001 Jasnow et al., 2006), with *one* demonstrating the role of testosterone on fear learning in pubescent mice (McDermott et al. 2012). Here we seek to determine the necessity of male and female circulating gonadal hormones at critical periods of hormone release toward the development of contextual fear learning. In doing so, we test the canonical “organizational hypothesis” that early perinatal and pubertal, but not later life circulating gonadal hormones are crucial for sex-specific maturation of context fear learning in male and female rats.

## Results and Discussion

In the *male rat*, the testes secrete testosterone, which peaks within hours of birth and continues to be released through the first few days of life before becoming quiescent (Rhoda et a., 1984; Corbier et al., 1992) until pubertal onset. Removal of testes by orchiectomy (oRX) within 8 hours of birth resulted in a reduction of context fear conditioning (Figure 1a) that was evident in adult but not in prepubertal juvenile male rats (Figure 1c: (Age x Surgery interaction F _1, 24_ = 4.761, p□<□.05; posthoc P24, *p* >.05, P90, *p* <.05). Given the absence of an effect during this prepubertal age, we tested whether androgens secreted during puberty were sufficient to produce the deficits evident during adulthood (Figure 1b). Elimination of circulating androgens by oRX days before puberty resulted in a similar reduction in adult context fear conditioning (*t* _(16)_ = 2.363, *p* < .05), indicating that preventing the pubertal surge of androgens was sufficient to disrupt adult context fear conditioning.

**Figure 1:**
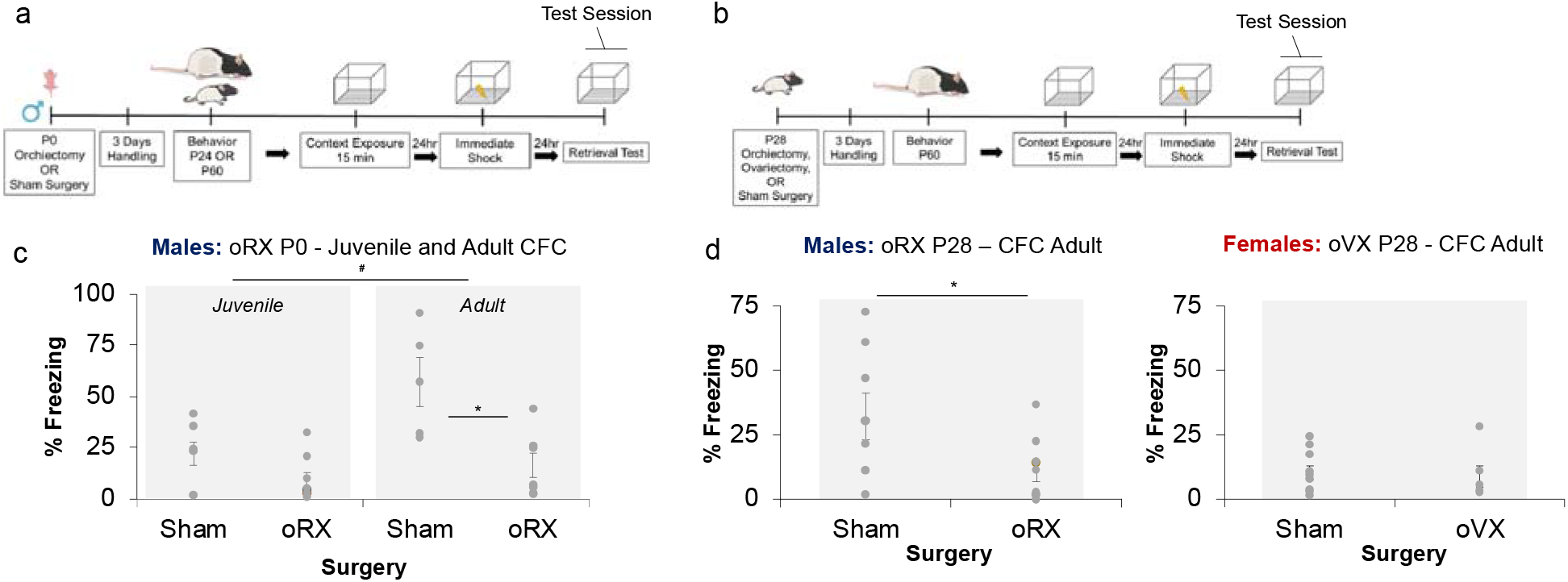
Deficits in Adult Context Fear Conditioning Following Neonatal and prepubertal oRX. Experimental timelines to examine the effects of (**a**) Neonatal orchiectomy on juvenile and adult CFC and (**b**) pre-pubertal orchiectomy and ovariectomy on adult CFC. (**c**) Removal of testes via oRX at birth significantly (* p < .05) reduced freezing in adults, but not in juvenile rats. Overall, conditioning was greater in adults than in juvenile rats (*p < .05). (**d**) oRX days before puberty similarly reduced CFC in adulthood (* p = .05), while removal of ovaries via ovariectomy (oVX) at birth did yield significant differences in conditioning.

In the *female rat*, while there is evidence that the ovaries secrete estrogens during infancy, this release is primarily described as latent until pubertal onset (Willing & Juraska, 2015). Thus, we initially tested the necessity of pubertal ovarian hormones by performing ovariectomies (oVX) in female rats days prior to puberty and tested adult context fear conditioning (Figure 1b). Unlike male rats, eliminating gonadal hormones at this developmental period did not significantly change adult context fear conditioning (Figure 1d) (*t* _(14)_ = .298, *p* > .05). However, since freezing performance in Sham and oVX females were both low, this may have limited the detection of group differences. To resolve this and determine if the male oRX-induced deficits in freezing resulted from a disruption of learning rather than an inability to freeze, in experiment 2, we included additional sessions of conditioning and testing (Figure 2a). Moreover, to determine if the adult introduction of gonadal hormones could rescue the effects of gonadectomy, we performed neonatal, prepubertal, and adult oRX and oVX and replaced circulating levels of testosterone (T) and 17β-estradiol (E) in adult male and female rats via subcutaneous implantation of silastic tubing (Yuasa et al.,1998; Kauffman et al., 2007, Zuloaga et al., 2011) days prior to conditioning. To confirm the efficacy of oRX, oVX and hormone replacement at the end of testing, we weighed male and female seminal vesicles and intrauterine horns (Tables 1 & 2).

**Table 1:** Seminal Vesicle Weight (Experiment 2)

|  | Male | Male | Male |
| --- | --- | --- | --- |
|  | Sham | oRX | oRX + T |
| P0-oRX → P60 harvest | 1.67 ± .12 | .35 ± .04* | .18 ± .01* |
| P28-oRX → P60 harvest | 1.45 ± .04 | .41 ± .14* | .97 ± .33 |
| P60-oRX → P90 harvest | 2.37 ± .10 | .26 ± .04* | 2.00 ± .5 |

\*Significantly differs from matched sham control group (p < .05)

**Table 2:** Uterine Horn Weight (Experiment 2)

|  | Female | Female | Female |
| --- | --- | --- | --- |
|  | Sham | oVX | oVX + E |
| P0-oVX → P60 harvest | 1.24 ± .28 | .35 ± .15* | .39 ± .08* |
| P28-oVX → P60 harvest | .75 ± .16 | .23 ± .12* | .64 ± .17 |
| P60-oVX → P90 harvest | .88 ± .25 | .66 ± .09* | .76 ± .30 |

\*Significantly differs from matched sham control group (p < .05)

**Figure 2:**
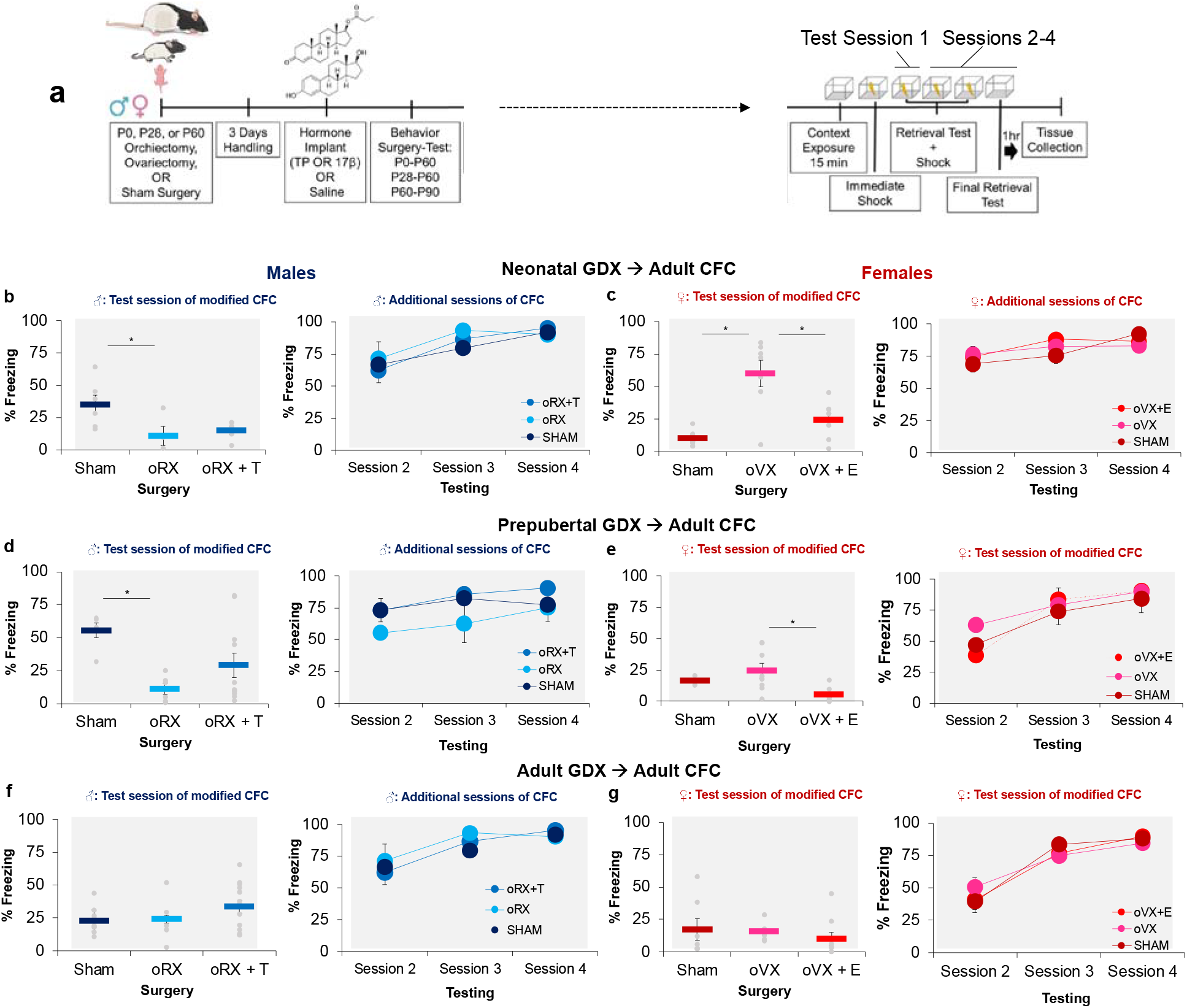
Sex-specific effects of gonadal removal during development upon CFC. Experimental timeline to examine the effects of orchiectomy or ovariectomy at birth, prior to puberty or adulthood and with or without prior hormone treatment upon adult CFC (**a**). (**b**) Neonatal oRX reduced adult CFC on test session 1 in comparison to sham males (* p < .05). (**c**) Neonatal oVX enhanced adult CFC on test session 1 in comparison to sham (**p < .01) and oVX + E females. (**d**) Prepubertal oRX reduced adult CFC on test session 1 in comparison to sham males (* p < .05). (**e**) Prepubertal oVX plus adult estrogen implantation reduced CFC on test sessions 1 and 2. (**f**) Adult oRX produced no significant effects. (**g**) Adult oVX produced no significant effects.

**Figure 3:**
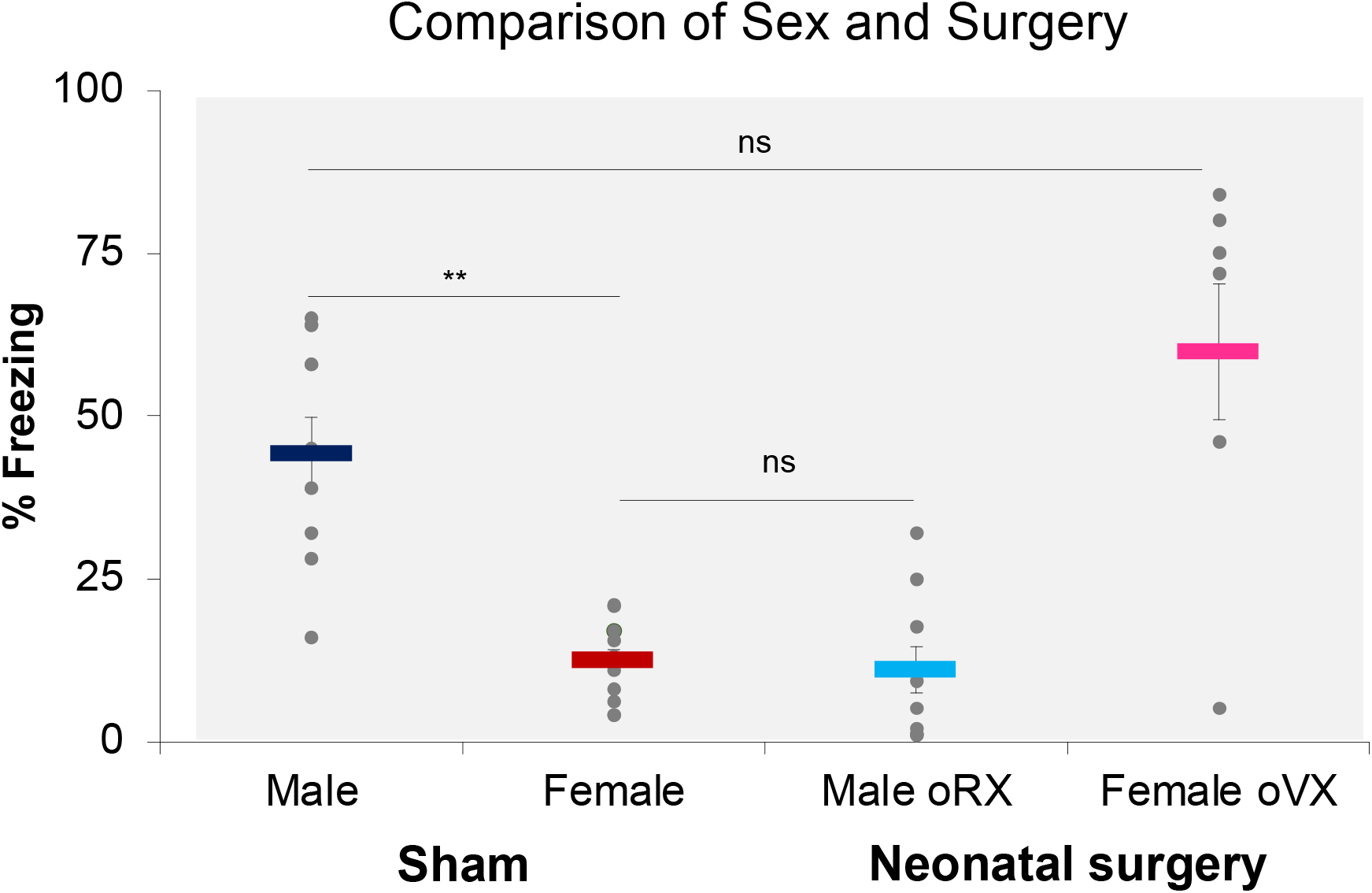
oVX resulted in male-like CFC and oRX resulted in female-like CFC. Context fear conditioning comparison between aggregate male and female sham with Neonatal oRX and oVX groups from experiment 2. Overall, sham males exhibited greater CFC than sham females (**p < .01), while similar levels of CFC were found between oVX and male sham rats and oRX (*ns*) and female sham rats (*ns*).

In experiment 2, once again in males, removal of the testes at birth resulted in a reduction in freezing after a single session of conditioning; however, with additional learning trials, this deficit was overcome, indicating that perinatal castration at best slowed the rate of learning (Figure 2b). This was supported by a significant interaction between Test sessions (1-4) and Condition (SHAM, oRX, oRX + T) (F _6, 39_ = 2.795, p□<□.05) and a follow-up between-subjects tests revealing a significant main effect of Condition on day 1 of testing (F _2, 13_ = 4.660, p□<□.05). Post-hoc analysis confirmed that oRX rats exhibited poorer conditioning than sham males (p < .05), which was not alleviated by T-replacement (p >.05). With additional sessions of training (2-4), there were no other differences in conditioning (day 2: F _2, 13_ = .180, p >.05; day 3: F _2, 13_ = 2.22, p >.05; day 4: F _2, 13_ = .523, p >.05), indicating multiple sessions of conditioning were required to mitigate the learning deficits in neonatal oRX males.

Accumulating evidence indicates that prepubertal estrogens are measurable within hours of birth and peak between 9 at 20 days postnatal (Döhler & Wuttke, 1975; Carson & Smith, 1986) and can contribute to the organization of female-specific reproductive-related behavior (Brock et al., 2011). We sought to determine whether this organizing role of ovarian hormones would extend to context fear conditioning. Indeed, in females that underwent oVX at birth, we observed an enhancement in context fear learning during adulthood that could be returned to sham levels with pre-training E treatment (Figure 2c). This finding was supported by a significant interaction between Test sessions and Condition (F _6, 60_ = 8.509, p□.<□.05) and follow-up between-subject tests revealing a significant main effect of Condition on day 1 of testing (F _2, 20_ = 15.848, p□<□.001) and confirmed by significant differences between Sham and oVX females (p < .01) and between oVX and oVX + E females (p <.01). Still, there were no significant differences between Sham and oVX + E females (p >.05), suggesting that the absence of early estrogens is permissive in the underlying organization of Context fear conditioning. With additional training, no other differences were detected between group conditions (day 2: F _2, 20_ = .326, p >.05; day 3: F _2, 20_ = 1.441, p > .05; day 4: F _2, 20_ = 1.347, p >.05).

As previously demonstrated in males, castration days prior to puberty reduced freezing behavior following the initial training session (Figure 2d). Moreover, as described in neonatal oRX rats, additional learning sessions could alleviate adult deficits in context fear conditioning. We tested whether androgens secreted during a pubertal window of development were sufficient to produce adult learning deficits and whether returning T before conditioning could rescue this deficit. Here we found that castration days before puberty significantly impeded the acquisition of contextual fear memories. This finding was supported by a significant interaction between Test sessions and Condition (F _6, 66_ = 3.103, p <.05). Follow-up between-subjects tests on the first test session revealed a significant main effect of Condition (F _2, 22_ = 5.173, p□<□.05) with oRX (p <.05) and oRX + T (p <.05) groups freezing significantly less than sham males. No group differences were found with additional training (day 2: F _2, 22_ = 2.395, p > .05; day 3: F _2, 22_ = 1.973, p > .05; day 4: F _2, 22_ = 2.212, p > .05).

In females, we confirmed that preventing that pubertal surge of estrogens did not alter context fear learning in adult rats. However, in prepubertal oVX rats, the adult introduction of E before conditioning reduced conditioning during the first session of training (Figure 2e). This was supported by a significant interaction between Test sessions and Conditions (F _6, 54_ = 3.205, p <.05) and a follow-up between-subject analysis of test session 1 (F _2, 16_ = 5.552, p < .05). Bonferroni post hoc analyses for Condition revealed that oVX alone females froze significantly more than oVX + E females (p < .05) on day 1. With no further differences in freezing between Sham and oVX females (p >.05). No differences were detected between groups on subsequent testing sessions (day 2: F _2, 16_ = 2.495 p >.05; day 3: F _2, 16_ = .503, p > .05; day 4: F _2, 16_ = .361, p > .05).

Prior work indicates that adult gonadectomies do not produce any effects on context fear conditioning (Anagnostaras et al.,1998; Gupta et al., 2001 McDermott et al., 2012). We confirmed that the adult removal of gonads in both males and females during early adulthood failed to produce any discernable effect on later context fear conditioning (Figure 2f-g), nor did hormone replacement in males (F _6, 60_ = .113, p□>□.05) or females (F _6, 36_ = .561, p□> □.05). There was a significant effect of Test Day in both males (F _3, 24_ = 98.058, p < □.01) and females (F _3, 17_ = 91.888, p < □.01), such that freezing increased across subsequent days of conditioning. These results confirm that the contributions of gonadal hormones toward context fear conditioning are specific to organizational periods of development that are evident with other sexually dimorphic motivated behaviors (McCullough et al., 1974; MacLusky & Naftolin,1981; Arnold & Breedlove, 1985; Lucion et al.,1996; Brock et al., 2011; Zuloaga et al, 2011; McDermott et al, 2012; Oyola & Handa, 2017).

Sex differences in context fear conditioning are well-established. While age- and litter-matched sham male and female rats were run in the first two studies of experiment 2, they were not powered to detect sex differences. Therefore, data from these studies were pooled and analyzed to evaluate potential sex differences and compare neonatal oRX and oVX neonatal rats during the 1st conditioning test session. This revealed a significant interaction between Sex and surgery Condition (F _1, 37_ = 60.892, p < .001). Post-hoc testing confirmed that sham males exhibited greater context fear conditioning than sham females (p < .001). Strikingly, conditioning in neonatal oVX females was comparable to sham males (p > .05) and neonatal oRX males showed similar learning to sham females (p > .05). Furthermore, the learning differences exhibited between sexes and oRX/oVX rats were not consistent with a deficit in reactivity to footshock (Sex: F _1, 35_ = .09, p > .05; oRX: F _1, 15_ = .142, p > .05; oVX: F _1, 17_ = 3.168, p >.05).

In summary, our data support the “organizational hypothesis” that gonadal hormones during the neonatal and peripubertal periods are vital to the sexual differentiation of context fear learning in male and female rats. To our knowledge, this is the first experimental evidence in males that the absence of neonatal and pubertal testicular hormones permanently disrupts the development of contextual fear conditioning. Perhaps more unprecedented is that the lack of ovarian hormones during the perinatal period enhanced the ability of females to acquire contextual fear memories while retaining the capacity to be modulated by initial exposure to estrogens during adulthood. This contrasted with the adult-specific removal of these hormones by oRX and oVX, where there were no disruptions in male or female contextual fear learning. Collectively, these results further support an organizational role of gonadal hormones in the sex differentiation of learning (Williams et al., 1990).

As discussed above, adult sex differences in context fear conditioning have been well-documented, with the current study lending further support for this. While estrogen in adult animals can modulate contextual-spatial-based learning, including context fear conditioning, we demonstrate that estrogens and androgens differentially organize this learning in male and female rats during development. It is important to note that neither oRX nor oVX altered reactivity to footshock unconditioned stimulus, suggesting that learning deficits result from a disruption contextual processing, associative learning, or both. At first glance, the effects of neonatal and prepubertal oRX suggest two organizational periods for the development of context fear conditioning. However, since the absence of gonadal hormones that spans infancy to adulthood and puberty to adulthood similarly produced an attenuation of context fear conditioning, this suggests that loss of gonadal hormones during puberty alone was sufficient to disrupt the organization of context fear conditioning and that loss of gonadal hormones at the start of infancy was not necessary to disturb the organization of this form of learning. This conclusion is further supported by experiment 1, which indicates that oRX performed at birth failed to disrupt prepubertal context fear conditioning. These results suggest that in males, the organization of context fear learning primarily occurs during puberty.

In female rats, the actions of gonadal hormones in the development and organization of context fear conditioning are more straightforward. The absence of gonadal hormones starting from infancy but not from puberty to adulthood enhanced learning. Indeed, counter to our initial prediction that gonadal hormones released during puberty represented a specific organizational window of action for maturation, our results indicate that estrogens produced and secreted by ovaries during infancy are organizational as found in reproductive-related behaviors (Brock et al., 2011). Strikingly, returning estrogen in adulthood in these lifelong postnatal gonadal hormone-deficient rats could attenuate this enhanced context fear learning. Overall, these results indicate that gonadal hormones during infancy have an organizational role in the maturation of context fear conditioning.

Collectively, we provide initial evidence that the organizational role of androgens and estrogens in context fear learning is uniquely timed to specific periods of development in male and female rats. In males, we provide evidence that androgens during puberty confer a continued maturation of context fear conditioning that is typical in males (Santarelli et al., 2018; Colón et al., 2018). In females, we demonstrate that estrogens are fundamental for developing female-typical patterns of contextual fear learning during the first few weeks of life. The translational relevance of these results may point toward the importance of circulating levels of estrogens during pre- and peri-natal periods that may predispose women to an enhanced capacity to learn and react to environmental threats evident in stress- and trauma-related disorders (Breslau et al., 1997; Kessler et al., 1994, Kessler et al., 1995; Kessler et al., 2005; Tolin & Foa, 2008; Breslau, 2009; McLean et al., 2011; Maeng & Milad, 2015).

## Materials and Methods

### Subjects

Subjects were male and female offspring of Long-Evans rats originally obtained from Envigo Laboratory and bred in the Life Sciences Research Building vivarium at the University at Albany. They were maintained on a 14:10-h light-dark cycle with free access to food and water. We designated postnatal (P) day 0 as the day of birth. Surgeries were performed on animals at either P0, P28, or P60. Rats were weaned at P21 and pair housed with same-sex littermates. All animal procedures followed the National Institutes of Health guidelines for the care and use of laboratory animals and were approved by the Institutional Animal Care and Use Committee of the University at Albany, SUNY.

### Surgery

#### Neonatal Orchiectomy

Pregnant dams were monitored daily for the appearance of pups. Within 8 hrs of birth, a total of male pups from different litters were removed from their homecage and anesthetized by hypothermia. Once pups were deeply anesthetized (on average 5 minutes), a single incision was made in the lower abdominal cavity to visualize the gonads. Animals were randomly assigned to either the sham or castrated groups. In the orchiectomy group, a ventral incision in the lower abdominal area was made through the skin and then the abdomen, after which testes were located on each side of the rat and removed. Sham surgery males received the same treatment for the same duration as castrated males, but the testes were left intact. Each surgery lasted an average of 10 minutes from induction to completion. A single suture was placed to close the abdominal cavity, and Vetbond glue was used to close the skin incision. Upon completion of the surgery, animals were placed on a heating pad and massaged until warm, and the righting reflex was restored. Once the animals were pink and moving, they were placed in a separate cage that contained bedding from their homecage. This was done to obscure the scent of the Vetbond glue, which dams appear to find aversive before the pups were brought back to their homepage (initial surgeries resulted in several pups not surviving the first post-surgical week due to rejection from the dam). All pups were removed from their mothers for a maximum of 2 hours. Post-surgical checks were performed daily for 7 days.

#### Prepubescent Orchiectomy and Ovariectomy

On P21, rats from separate breeding pairs were weaned from their mothers and pair housed by sex. On P28, males and females were removed from the vivarium for surgery. Using 2.5% isoflurane gas anesthesia, each gonadectomy and sham surgery was completed with an average duration of 25 minutes from induction to completion. Both ovariectomies and orchiectomies were performed with a ventral incision in the lower abdominal area through the skin and then another through the muscle wall, after which testes or ovaries were located on each side of the rat, ligated with absorbable surgical thread, and removed. The muscle incision was closed using absorbable sutures, and the skin incision was closed using both suture and Vetbond surgical glue. Animals were then placed in a separate cage until they fully recovered from anesthesia before being placed back with their littermate. Identical skin and abdominal incisions were made on sham animals, but the gonads remained intact; the duration of isoflurane exposure was equal to that of castrated animals. Rats were returned to the vivarium after the restoration of the righting reflex. Following surgery, all rats were administered Carprofen (1mg/kg) subcutaneously for 2 consecutive days and monitored for 7 days.

#### Adult Orchiectomy and Ovariectomy

P60 male and female rats were removed from the vivarium for surgery. Surgical castration and sham procedures were identical to the procedures described in prepubescent animals.

#### Hormone replacement

Five days before the start of behavior testing adult animals underwent a brief surgery (approximately 8 minutes) to implant Silastic tubing subcutaneously. The Silastic tubes were filled with either testosterone propionate (T) or 17-beta estradiol (E) or remained empty and sealed at both ends with medical-grade silicone adhesive (Dow Corning Corp., Midland, MI). All sham surgery animals received empty Silastic capsules. Castrated animals were divided into GDX/OVX + hormone capsule or GDX/OVX + empty capsule. Male castrated rats received two Silastic (Dow Corning Corp. Midland, MI) capsules (1.57 mm inner diameter, 3.18 mm outer diameter; 20 mm effective release length) containing testosterone propionate or 2 empty capsules of the same length and diameter via a 2-cm incision dorsally at the nape. Female castrated rats received a Silastic (Dow Corning Corp., Midland, MI) capsule (1.27 mm inner diameter, 3.18 mm outer diameter; 20 mm effective release length) containing 17-beta estradiol or an empty capsule of the exact dimensions via a 2-cm incision dorsally at the nape. It has been demonstrated that the amount of respective hormone (E or T) delivered through the Silastic capsules results in circulating levels of the hormone that are consistent with normal physiological levels of circulating hormone in intact adult rats (Kauffman et al., 2007; Zuloaga et al., 2011).

### Behavioral Testing

#### Apparatus

##### Context Fear Conditioning and Testing Experiment 1

All rats were handled for two minutes each on three consecutive days before behavior testing. Across all behavior experiments (1-3), we used a context pre-exposure facilitation procedure described in Colon et al. (2018) and Colon & Poulos (2020). In experiment 1, male P24 (juvenile), P60, or P90 (adult) rats that previously underwent castration or sham surgery at P0 were pre-exposed to the conditioning chamber for 15 min. Twenty-four hours later, the animals were brought back to the conditioning chamber. After 10 seconds in the chamber, rats received a footshock (1.0 mA; 2-sec duration) and were removed immediately from the chamber. Twenty-four hours later, rats were returned to the conditioning chamber for a 4-minute context fear test.

##### Experiment 2

Based on the results of experiment 1, we were interested in whether additional learning trials could rescue low freezing. On day 1, animals were pre-exposed to the context for 15 minutes. On day 2, they were placed back into the context and received an immediate footshock (1mA, 2 sec) 10s after being placed into the chamber. On days 3-5, animals were returned to the conditioning chamber for a 4-minute retrieval test followed by a footshock at the end of the test session (total time spent in context was 242 sec), which was repeated for two additional sessions. On day 6, the final retrieval test session, there was no footshock presented.

##### Perfusion and Organ collection

Animals were administered Euthanasia-III Solution (sodium pentobarbital) and transcardially perfused with 1% potassium phosphate buffer solution and 4% paraformaldehyde (Sigma-Aldrich; MO, USA). The seminal vesicles of male subjects and the uterine horns of female subjects were removed and weighed. In males, the seminal vesicles (reproductive glands) are sensitive to circulating androgens, and organ weights can be used as indicators of androgen status in rats and mice (Champin & Creasy, 2012); similarly, uterine horn width and weight are positively correlated with circulating concentrations of estrogens, particularly 17-beta estradiol, in mice and rats and can also be used as an indicator of circulating estrogens (Wood et al., 2007).

## References

Anagnostaras SG, Maren S, DeCola JP, Lane NI, Gale GD, Schlinger BA, Fanselow MS. Testicular hormones do not regulate sexually dimorphic Pavlovian fear conditioning or perforant-path long-term potentiation in adult male rats. Behav Brain Res. 1998 Apr;92(1):1–9. doi: 10.1016/s0166-4328(97)00115-0. PMID: 9588680.

Arnold AP. The organizational-activational hypothesis as the foundation for a unified theory of sexual differentiation of all mammalian tissues. Horm Behav. 2009 May;55(5):570–8. doi: 10.1016/j.yhbeh.2009.03.011. PMID: 19446073; PMCID: PMC3671905.

Arnold AP, Breedlove SM. Organizational and activational effects of sex steroids on brain and behavior: a reanalysis. Horm Behav. 1985 Dec;19(4):469–98. doi: 10.1016/0018-506x(85)90042-x. PMID: 3910535.

Brock O, Baum MJ, Bakker J. The development of female sexual behavior requires prepubertal estradiol. J Neurosci. 2011 Apr 13;31(15):5574–8. doi: 10.1523/JNEUROSCI.0209-11.2011. PMID: 21490197; PMCID: PMC3085119.

Carson R, Smith J. Development and steroidogenic activity of preantral follicles in the neonatal rat ovary. J Endocrinol. 1986 Jul;110(1):87–92. doi: 10.1677/joe.0.1100087. PMID: 3090186.

Colon L, Odynocki N, Santarelli A, Poulos AM. Sexual differentiation of contextual fear responses. Learn Mem. 2018 Apr 16;25(5):230–240. doi: 10.1101/lm.047159.117. PMID: 29661835; PMCID: PMC5903402.

Corbier P, Edwards DA, Roffi J. The neonatal testosterone surge: a comparative study. Arch Int Physiol Biochim Biophys. 1992 Mar-Apr;100(2):127–31. doi: 10.3109/13813459209035274. PMID: 1379488.

Daniel JM, Dohanich GP. Acetylcholine mediates the estrogen-induced increase in NMDA receptor binding in CA1 of the hippocampus and the associated improvement in working memory. J Neurosci. 2001 Sep 1;21(17):6949–56. doi: 10.1523/JNEUROSCI.21-17-06949.2001. PMID: 11517282; PMCID: PMC6763069.

Dohanich, G. (2002). Gonadal steroids, learning, and memory. Hormones, brain and behavior, 265–327.

Döhler KD, Wuttke W. Changes with age in levels of serum gonadotropins, prolactin and gonadal steroids in prepubertal male and female rats. Endocrinology. 1975 Oct;97(4):898–907. doi: 10.1210/endo-97-4-898. PMID: 1193012.

Gupta RR, Sen S, Diepenhorst LL, Rudick CN, Maren S. Estrogen modulates sexually dimorphic contextual fear conditioning and hippocampal long-term potentiation (LTP) in rats(1). Brain Res. 2001 Jan 12;888(2):356–365. doi: 10.1016/s0006-8993(00)03116-4. PMID: 11150498.

Jasnow AM, Schulkin J, Pfaff DW. Estrogen facilitates fear conditioning and increases corticotropin-releasing hormone mRNA expression in the central amygdala in female mice. Horm Behav. 2006 Feb;49(2):197–205. doi: 10.1016/j.yhbeh.2005.06.005. Epub 2005 Aug 3. PMID: 16083887.

Kauffman AS, Gottsch ML, Roa J, Byquist AC, Crown A, Clifton DK, Hoffman GE, Steiner RA, Tena-Sempere M. Sexual differentiation of Kiss1 gene expression in the brain of the rat. Endocrinology. 2007 Apr;148(4):1774–83. doi: 10.1210/en.2006-1540. Epub 2007 Jan 4. PMID: 17204549.

Korol DL. Role of estrogen in balancing contributions from multiple memory systems. Neurobiol Learn Mem. 2004 Nov;82(3):309–23. doi: 10.1016/j.nlm.2004.07.006. PMID: 15464412.

Lenz KM, McCarthy MM. Organized for sex -steroid hormones and the developing hypothalamus. Eur J Neurosci. 2010 Dec;32(12):2096–104. doi: 10.1111/j.1460-9568.2010.07511.x. PMID: 21143664; PMCID: PMC5350613.

Luine VN. Sex steroids and cognitive function. J Neuroendocrinol. 2008 Jun;20(6):866–72. doi: 10.1111/j.1365-2826.2008.01710.x. Epub 2008 Jun 1. PMID: 18513207.

MacLusky NJ, Naftolin F. Sexual differentiation of the central nervous system. Science. 1981 Mar 20;211(4488):1294–302. doi: 10.1126/science.6163211. PMID: 6163211.

McCullough, J., Quadagno, D. M., & Goldman, B. D. (1974). Neonatal gonadal hormones: Effect on maternal and sexual behavior in the male rat. Physiology & Behavior, 12(2), 183–188.

McDermott CM, Liu D, Schrader LA. Role of gonadal hormones in anxiety and fear memory formation and inhibition in male mice. Physiol Behav. 2012 Mar 20;105(5):1168–74. doi: 10.1016/j.physbeh.2011.12.016. Epub 2011 Dec 28. PMID: 22226989.

Morgan MA, Pfaff DW. Effects of estrogen on activity and fear-related behaviors in mice. Horm Behav. 2001 Dec;40(4):472–82. doi: 10.1006/hbeh.2001.1716. PMID: 11716576.

Oyola MG, Handa RJ. Hypothalamic-pituitary-adrenal and hypothalamic-pituitary-gonadal axes: sex differences in regulation of stress responsivity. Stress. 2017 Sep;20(5):476–494. doi: 10.1080/10253890.2017.1369523. Epub 2017 Aug 31. PMID: 28859530; PMCID: PMC5815295.

Rhoda J, Corbier P, Roffi J. Gonadal steroid concentrations in serum and hypothalamus of the rat at birth: aromatization of testosterone to 17 beta-estradiol. Endocrinology. 1984 May;114(5):1754–60. doi: 10.1210/endo-114-5-1754. PMID: 6714163.

Schulz KM, Sisk CL. The organizing actions of adolescent gonadal steroid hormones on brain and behavioral development. Neurosci Biobehav Rev. 2016 Nov;70:148–158. doi: 10.1016/j.neubiorev.2016.07.036. Epub 2016 Aug 4. PMID: 27497718; PMCID: PMC5074860.

Taxier LR, Gross KS, Frick KM. Oestradiol as a neuromodulator of learning and memory. Nat Rev Neurosci. 2020 Oct;21(10):535–550. doi: 10.1038/s41583-020-0362-7. Epub 2020 Sep 2. PMID: 32879508; PMCID: PMC8302223.

Williams CL, Barnett AM, Meck WH. Organizational effects of early gonadal secretions on sexual differentiation in spatial memory. Behav Neurosci. 1990 Feb;104(1):84–97. doi: 10.1037//0735-7044.104.1.84. PMID: 2317288.

Willing J, Juraska JM. The timing of neuronal loss across adolescence in the medial prefrontal cortex of male and female rats. Neuroscience. 2015 Aug 20;301:268-75. doi: 10.1016/j.neuroscience.2015.05.073. Epub 2015 Jun 3. PMID: 26047728; PMCID: PMC4504753.

Wood GA, Fata JE, Watson KL, Khokha R. Circulating hormones and estrous stage predict cellular and stromal remodeling in murine uterus. Reproduction. 2007 May;133(5):1035–44. doi: 10.1530/REP-06-0302. PMID: 17616732.

Yuasa H, Ono Y, Fukabori Y, Ohma C, Yamanaka C, Suzuki K. Usefulness of the estrogen releasing silastic tubing, The Kitakanto Medical Journal, 1998, Volume 48, Issue 1, Pages 15–18, Released on J-STAGE October 21, 2009, Online ISSN 1881-1191, Print ISSN 1343-2826, https://doi.org/10.2974/kmj.48.1

Zuloaga DG, Jordan CL, Breedlove SM. The organizational role of testicular hormones and the androgen receptor in anxiety-related behaviors and sensorimotor gating in rats. Endocrinology. 2011 Apr;152(4):1572–81. doi: 10.1210/en.2010-1016. Epub 2011 Feb 15. PMID: 21325044; PMCID: PMC3060630.

